# Building an Ecosystem of Seizure Localization Methods: Neural Fragility as the First Step

**DOI:** 10.1101/2025.09.11.675707

**Authors:** Jiefei Wang, Anne-Cecile Lesage, Oliver Zhou, Ioannis Malagaris, Sean O'Leary, Liliana Camarillo Rodriguez, Diosely C Silveira, Zhengjia Wang, Yuanyi Zhang, Patrick J Karas

## Abstract

The current treatment for drug-resistant epilepsy (DRE) is surgical intervention, which relies on accurate identification of the seizure onset zone (SOZ) using intracranial EEG (iEEG) data. iEEG analysis with computational epileptogenic zone identification algorithms (CEZIAs) is a promising step towards better SOZ localization and surgical outcomes. A key step in validation and adoption of CEZIAs is to allow for widespread shared development and validation of code and data. We describe a set of three R packages to achieve this goal. Our ecosystem of seizure localization methods involves a straightforward analysis pipeline, standardized data formatting and storage, and completely documented and open-source code. The TableContainer package allows for easy storage and manipulation of table data, serving as groundwork for the Epoch package, which is specifically geared towards iEEG data. The Epoch package allows for cropping, resampling, and visualization of iEEG data and provides publicly downloadable iEEG data for reproducibility. Finally, the EZFragility package uses these two foundational packages to analyze iEEGs for SOZ localization using the Neural Fragility method described by Li et al. EZFragility was built using the same core principles as the original method but included several enhancements in computational efficiency and user experience. It accurately reproduces neural fragility results for both sample patients used in the original paper. This project serves as the first step towards building an open-source, reproducible ecosystem of seizure localization methods in R. Future steps include the addition of other CEZIAs using the framework and sample data already made available by these packages.

**Significance Statement:** Localization of the seizure onset zone (SOZ) is a critical step in surgical treatment of drug-resistant epilepsy. Computational epileptogenic zone identification algorithms (CEZIAs) are promising potential tools to aid in clinical decision-making. However, shared development and verification of CEZIAs is difficult due to obscured source code, variable data structures, and limited data availability. The EZFragility software package^1^ is the first step in building a collaborative ecosystem of CEZIAs that can be downloaded, tested, and used without these roadblocks. EZFragility^1^ and its dependent packages TableContainer^2^ and Epoch^3^ are freely available on the Comprehensive R Archive Network (CRAN), with source code viewable on GitHub. They provide an open-source framework for CEZIA code, data formatting, and data access, all with extensive documentation.

## Introduction

Drug-resistant epilepsy (DRE) affects approximately 15 million individuals worldwide^4,5^ and is defined by persistent seizures despite adequate trials of two anti-seizure medications. It has profound psychosocial and economic impacts^6,7^ and increases risk of premature mortality^8^. For patients with focal DRE, surgical resection of the seizure onset zone (SOZ), the brain region critical for seizure generation and propagation, remains the most effective treatment^9,10^.

Localization of the SOZ is a complex process. The current clinical standard relies on expert epileptologist interpretation of seizures recorded via intracranial electroencephalography (iEEG)^11^. Epileptologists visually examine seizure onset patterns and epileptiform discharges across implanted electrodes, integrating this information with anatomical and clinical data to identify the SOZ. However, even with invasive monitoring, postoperative seizure freedom rates range from only 30% to 70%^12–14^. This variability underlines the challenges of SOZ localization and the need for more objective, reproducible methods.

Computational epileptogenic zone identification algorithms (CEZIAs) have emerged as promising assistants to clinical decision-making. CEZIAs extract subsets of features from iEEG recordings, often focusing on the temporal dynamics, frequency signatures, and anatomical distribution of ictal signals. These features represent specific “anatomical-electro-clinical correlations” fundamental to defining the SOZ^15^. CEZIAs vary in complexity, from univariate ictal analyses of individual channels to multivariate ictal assessments of large-scale brain networks. Though promising, widespread clinical adoption and validation of CEZIAs remains limited by the following critical barriers:

1. **Code Accessibility**: CEZIA source code is often unavailable, outdated, or written in proprietary programming languages^15^. Many CEZIA analyses are performed with in-house code and limited documentation, hindering independent reproduction. For example, the neural fragility^16,17^ methodology does not include source code or complete specification of implementation parameters.
2. **Data Structure Standardization**: There is no standardized data structure in R for the iEEG matrix and critical clinical information (e.g., electrode location, surgical outcomes). Though there is the iEEG-BIDS format for raw data^18^, essential analysis information such as patient demographics, channel information, and other metadata is dispersed across individual files in various folders. Researchers are forced to consolidate these key elements for their analysis, hindering the reproducibility and comparability of CEZIAs across studies.
3. **Data Availability**: While some public iEEG datasets exist (e.g. HUP^19^, Fragility^20^), they require substantial preprocessing and may have faulty electrodes or poor implantations that do not properly capture the SOZ. It is difficult to find properly formatted, high-quality data ready for CEZIA analysis.

Overcoming these challenges requires not just new algorithms but a unified, open-source computational ecosystem for CEZIA development, validation, and deployment. We describe a set of R packages to achieve this goal, creating an ecosystem for the standardized implementation of computation seizure localization methods including straightforward analysis pipelines, standardized data formatting and storage, and completely documented and open-source code. We implemented neural fragility within this ecosystem as the first CEZIA. The neural fragility algorithm represents the brain as an epileptogenic network and was shown to outperform 20 other iEEG biomarkers in predictive power and interpretability^13^.

In this manuscript, we describe our complete open-science implementation of neural fragility in R, incorporating open iEEG data, signal processing tools, packages for data format standardization, and open-source code for visualization. This work represents a significant step toward establishing a reproducible and sustainable framework for CEZIAs in epilepsy research.

## Materials and Methods

### Pipeline

Going from raw data to CEZIA analyses involves several key steps. Figure 1 represent the conceptual workflow for CEZIA analysis. First, the data must be acquired and preprocessed. Preprocessing was performed with RAVE (Reproducible Analysis & Visualization of iEEG), an open-source, NIH-funded software built in R^21^. Data format was then standardized using the TableContainer and Epoch packages. Last, the standardized data will be analyzed and visualized by the EZFragility package. To simplify the workflow, we uploaded the standardized data for the fragility analysis to the Open Science Framework^22^, allowing future users using this data to skip directly to the last step, CEZIA analysis and visualization. Our packages are built upon popular R packages such as ggplot2, foreach, and Shiny^23^ to maximize usability and flexibility.

**Figure 1.**
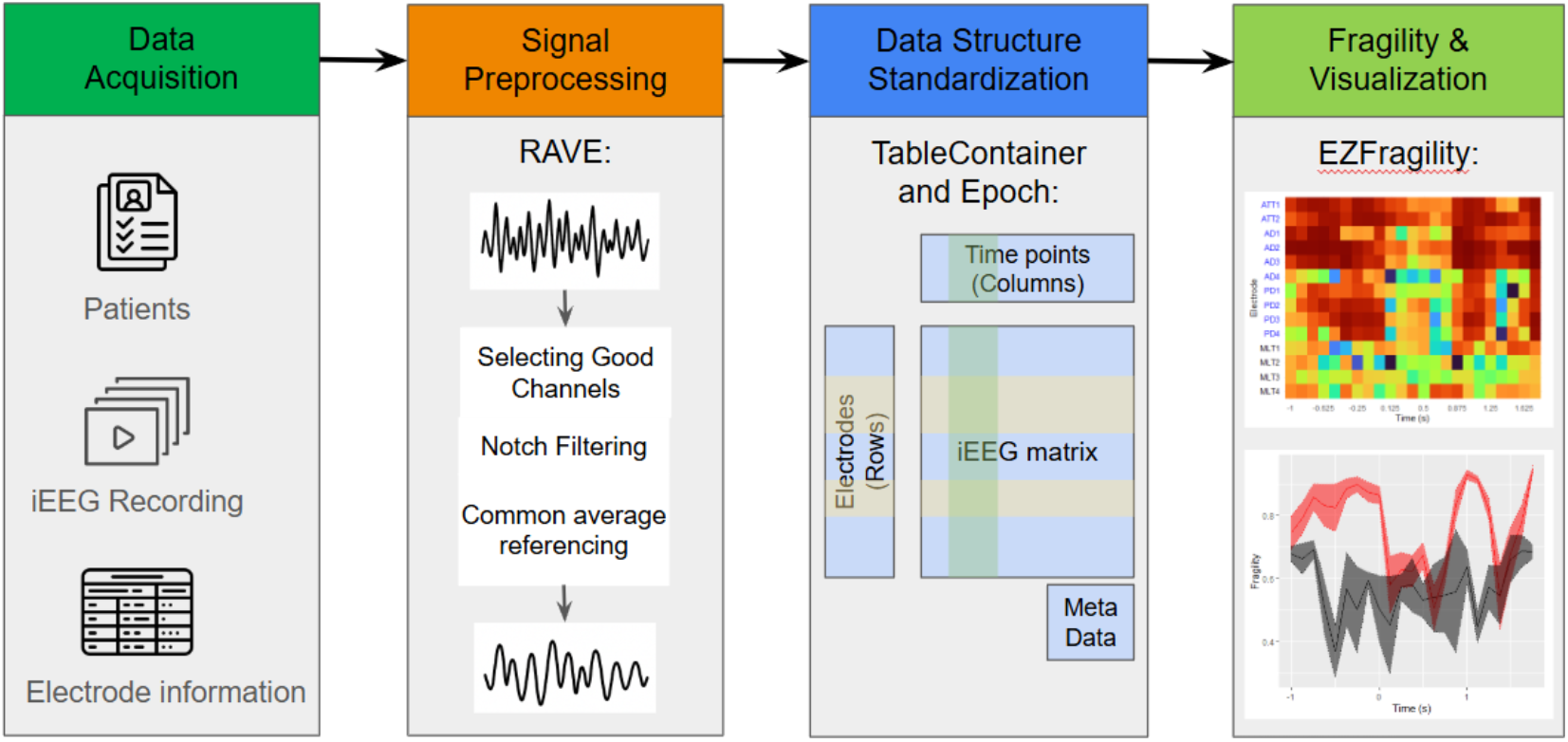
provides a conceptual workflow for SOZ prediction using the R packages RAVE, TableContainer, Epoch, and EZFragility.

### Data acquisition and preprocessing

The first step in the pipeline involves acquiring and preprocessing iEEG data suitable for fragility analysis. The seizure iEEG dataset from the Fragility multi-center retrospective study on OpenNeuro was used for this implementation^20^. This dataset choice provides clinically validated seizure recordings with known outcomes as well as enables direct validation of the EZFragility package implementation against the original Fragility method results. All of the following steps were performed using RAVE^21^. The imported data was Notch filtered for contamination identified upon visual inspection, commonly on harmonics of 60 Hz (60, 120, 180, etc.), with cutoff windows of 1, 2, and 2 Hz, respectively. If not already excluded during data import, faulty channels were removed by visual inspection. We excluded those that were clinically determined to be excessively noisy, those recording from white matter, and other non-EEG electrodes (e.g., EKG). Next, a common average reference across the remaining electrodes was used to remove any correlated noise. Finally, an epoch was generated, specifying the seizure onset times and other notable events for each recording. The time windows for analysis are chosen relative to the seizure onset times (e.g., 30 seconds before onset to 30 seconds after), which are specified in the epoch. Voltage data for this time window was extracted, resulting in an M-by-N matrix of voltage values over the time window where M is the number of electrodes and N is the number of time points. All data preprocessing steps, including channel quality assessment, artifact identification, and seizure onset marking, were performed under the supervision of a senior clinical epileptologist (DCS) to ensure clinical accuracy and validity.

### Data Structure Standardization

The TableContainer^2^ and Epoch^3^ R packages were developed as a data container to establish a standardized data structure for iEEG analysis. The TableContainer^2^ package provides a flexible framework for storing tabular data, the row and column metadata, and the metadata for the table as a whole. Subsetting the data will also subset the metadata, ensuring that the metadata remains consistent with the data.

The Epoch package extends TableContainer^2^ to specifically handle iEEG data, allowing for easy manipulation (e.g., cropping and resampling) and visualization of iEEG recordings. The main function is *Epoch*, which creates an Epoch object from the iEEG data. The Epoch object contains the iEEG recordings, electrode information as row metadata (e.g., name, labeled as in the seizure onset zone (SOZ) or not, labeled as in the resection area or not), and patient information as metadata (e.g., gender, surgical outcome, brain region hypothesis, age at first seizure, age at iEEG epilepsy surgery evaluation, type of electrodes (ECOG and SEEG)). This way, the Epoch object can provide a complete representation of the seizure iEEG data for downstream analysis.

Furthermore, we have curated the Epoch data obtained from the Fragility multi-center retrospective study and made them publicly available on the Open Science Framework (OSF)^22^. We provide the *EpochDownloader* function in the Epoch package, which can directly download the curated iEEG data from R. *EpochDownloader* allows other researchers to access the standardized iEEG data used in the EZFragility package and reproduce our results in this manuscript using the same data.

### Fragility calculation

The EZFragility^1^ package implements the neural fragility algorithm as described in Li et al^17^. The fragility metric quantifies the minimum perturbation at a single region required to drive the iEEG network from a stable state to an unstable state. The fragility values are normalized using a reverse scaling, with higher fragility values indicating regions more prone to introducing unstable neural dynamics. The core function, *calcAdjFrag*, takes an Epoch object, window size, and step size as input. It computes the fragility vector within a time window of the Epoch iEEG matrix. Within a window, the function constructs a linear model with LASSO penalty for modeling the neural dynamics and calculates the minimum two-norm perturbation matrix required to destabilize the system with an eigenvalue eig=σ+iω with a norm of one. The fragility value for each electrode is defined as the minimum perturbation norm across all searched ω. Finally, the fragility values from electrodes are pooled together and normalized using a reverse scaling: (max_fragility - fragility_value) /max_fragility. This forms the fragility vector for each time window. The algorithm repeats the above calculation while sliding the time window by a step size unit. The final result is a fragility matrix, with each column being a fragility vector from a time window. The value of the fragility matrix ranges from 0 to 1, where higher values indicate greater fragility. This normalization allows for a direct comparison of fragility values across different electrodes and time windows.

We developed a custom algorithm to automatically determine the LASSO penalty used in the algorithm, which was not clearly specified in Li et al^13^. The algorithm employs an adaptive regularization parameter (lambda) selection strategy through binary search when no lambda value is specified by the user. The *ridgeSearch* function first attempts to fit the linear model with a small initial lambda value (1e-4) as the original author used. If the resulting dynamics are unstable (indicating insufficient regularization), the algorithm automatically searches for an optimal lambda within the range [1e-4, 10] using binary search over up to 20 iterations until the neural dynamics become stable. The user can opt out of this behavior by providing the desired penalty strength. However, it should be noted that the behavior of the fragility matrix from an unstable neural dynamic is ill-defined.

Additional efforts have been made to enhance computational efficiency and user experience. We implemented a custom ridge regression algorithm specifically tailored for the fragility calculation. The ridge regression model strips unnecessary calculations important for statistical analysis but not for the fragility model. It is able to gain over 10x performance improvement compared to glmnet::glmnet. Furthermore, the package supports parallel computing through the foreach package, enabling distributed computation across multiple CPU cores to significantly reduce processing time for large datasets. A progress bar provides real-time feedback during computation, which is particularly useful for impatient users given the computationally intensive nature of the fragility calculations.

The output of *calcAdjFrag* is a Fragility object that contains the fragility matrix and its metadata. The function *estimateSOZ* can be used to estimate the seizure onset zone (SOZ) based on the Fragility object. It aggregates the fragility values across time windows for each electrode using a user-specified method (mean, median, max, or min) and ranks the electrodes based on their aggregated fragility scores. The function returns the indices or names of the top N electrodes with the highest fragility scores, where N is determined by a user-specified proportion of the total electrode count (default 10%). These electrodes are considered most likely to be critical in seizure generation and propagation. This provides a practical and interpretable cue for clinicians and researchers to further investigate those identified electrodes.

### Visualization

We developed a set of visualization functions within the EZFragility^1^ package to facilitate the interpretation of fragility analysis results. The visualization functions take a Fragility object (the output of *calcAdjFrag*) as input and generate a ggplot^24^ object that can be further customized by users. The plotting framework implemented in this function reproduces the methods used in the neural fragility original paper to visualize correlations between fragility and SOZ electrodes.

The *plotFragHeatmap* function generates a spatiotemporal heatmap of the fragility matrix (see Figure 2 in the results section), with electrodes displayed along the y-axis and time windows along the x-axis. Color intensity represents fragility values, with warmer colors indicating higher fragility and potential seizure onset zones. The function automatically scales colors to highlight regional differences. Users can optionally group electrodes to highlight them in the heatmap. This visualization provides an intuitive overview of fragility dynamics throughout the seizure recording.

**Fig 2.**
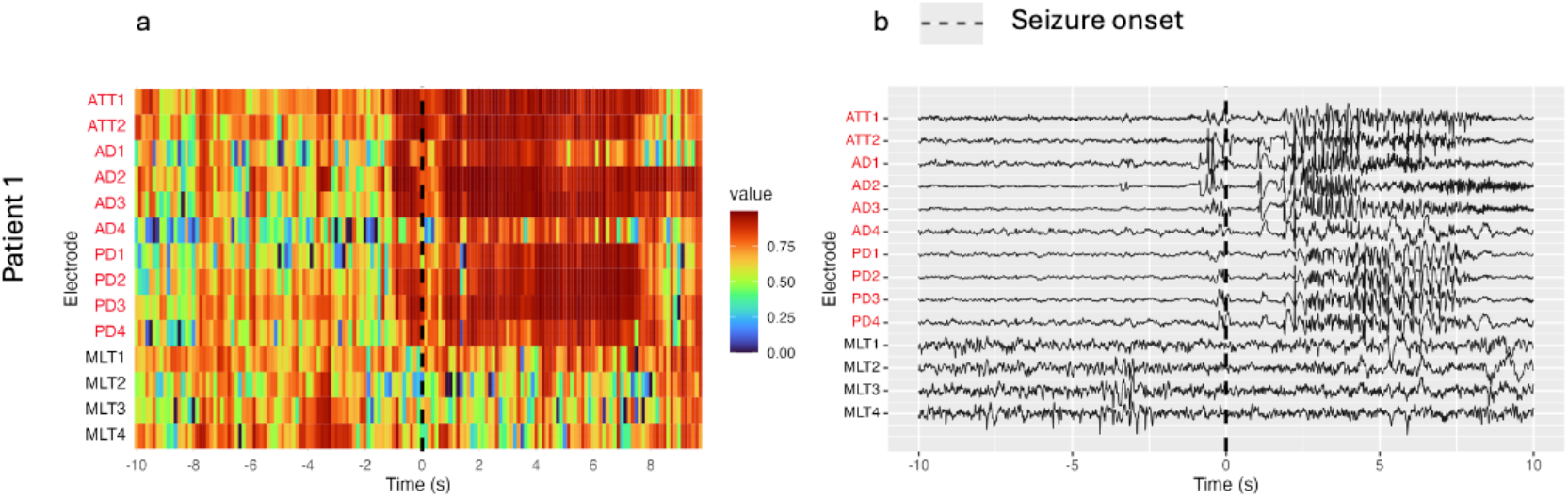
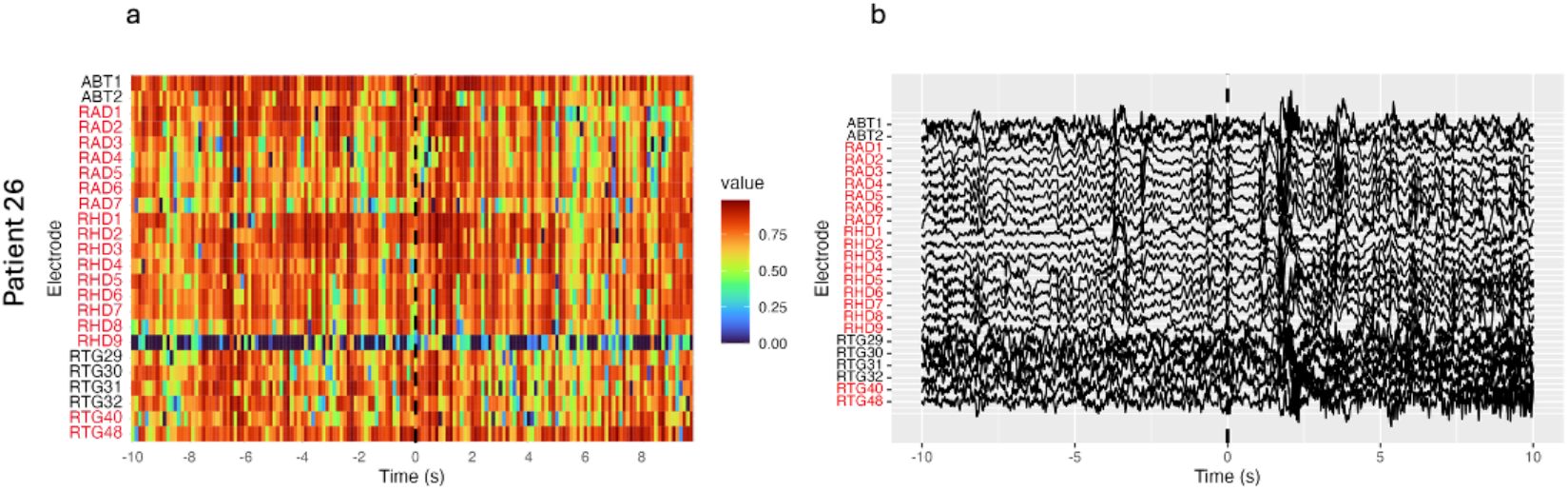
Fragility heatmap results (a) with raw voltage plot (b) for patient 01 (success) and patient 26 (failure) produced with the EZFragility R package. Electrodes highlighted in red were identified as SOZ by clinical epileptologists. On the fragility map, values closer to 1 are more fragile. Note the correlation between fragile regions and identified SOZ electrodes in PT01 (surgical success), and the lack thereof for PT26 (surgical failure).

To compare fragility across different groups of electrodes (e.g., seizure onset zone vs. non-seizure onset zone), EZFragility^1^ provides the *plotFragDistribution* and *plotFragQuantile* functions. The *plotFragDistribution* function creates line plots for each group, showing mean fragility values across time windows with shaded areas representing standard deviation or standard error. Similarly, the *plotFragQuantile* function displays fragility values as quantiles for each time window and compares them across two groups. These functions allow researchers to visually assess differences in fragility patterns between groups and help identify time windows where distributional differences are most pronounced.

## Code availability

Packages TableContainer^2^, Epoch^3^, and EZFragility^1^ will be available in the CRAN library. The GitHub packages with installation instructions, readme, and example vignettes are accessible through the following links:

- TableContainer: https://github.com/Jiefei-Wang/TableContainer
- Epoch: https://github.com/Jiefei-Wang/Epoch
- EZFragility: https://github.com/Jiefei-Wang/EZFragility

The script for reproducing the results shown in the manuscript is the vignette manuscript_reproducible.Rmd within the package EZFragility.

## Results

The EZFragility package can reproduce results from Li et al. Two out of the three patients used as examples for Figure 4 in the 2021 neural Fragility paper^17^ were made publicly available in the Fragility dataset. Of these patients, PT01 was a surgical success, with the high-fragility electrodes identified by the algorithm matching those identified by clinical epileptologists. In contrast, PT26 was a surgical failure, and the high-fragility electrodes were not consistent with clinical EEG interpretation. The raw data was band-pass-filtered between 0.5 and the Nyquist frequency with a fourth-order Butterworth filter. The vignette manuscript_reproducible.Rmd in the EZFragility package describes how to obtain the data and reproduce the validation figures of this paper for both patients.

Figure 2 shows the voltage plots and fragility maps generated by the EZFragility package for the first recorded seizure of PT01 and PT26, 10 seconds before and after seizure onset. Our findings were consistent with the original Fragility paper, both in terms of high fragility electrode identification and visual inspection. Furthermore, analysis of average fragility over time between clinically identified SOZ electrodes and non-SOZ electrodes was also consistent with the original paper (Figure 3). This figure reproduces the extended figure 4 results of the Li et al. original paper^17^.

**Fig. 3.**
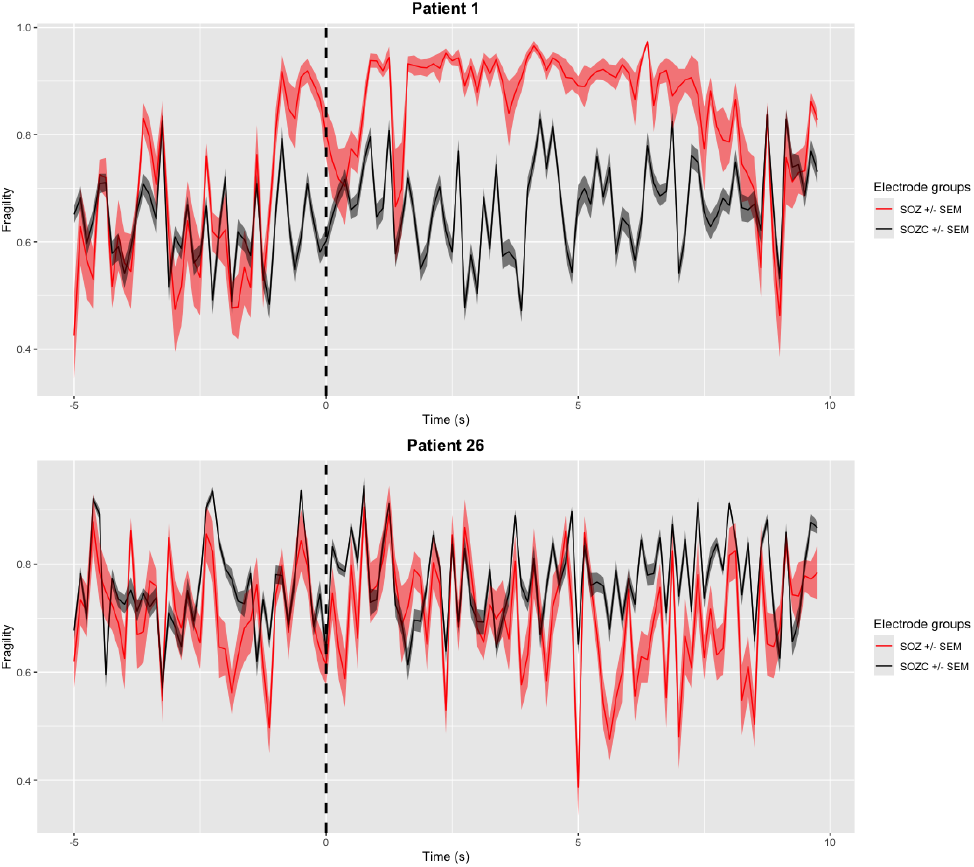
Analysis of average fragility over time between clinically identified SOZ electrodes and non-SOZ electrodes for patient 01 (success, top) and patient 26 (failure, bottom). Note how the clinically identified SOZ electrodes have higher average fragility after seizure onset for patient 01 while there is no significant difference for patient 26.

## Discussion

The TableContainer^2^, Epoch^3^ and EZFragility^1^ packages serve as the first implementation in a suite of R packages designed to build a comprehensive R ecosystem for CEZIA development and deployment. This system of packages includes a framework for data structure and manipulation, ready-to-use sample data, and rigorous but efficient computational algorithms. Additional CEZIAs will be easier to implement with this structure and workflow as a foundation. Along with RAVE, an established R platform for iEEG analysis and visualization, these packages serve as an important step towards an open-science, collaborative environment of computation iEEG analysis.

To address the problem of code accessibility, the packages and documentation for the functions within them will all be publicly available on GitHub as well as the Comprehensive R Archive Network. This will allow users to not only inspect and work with the code themselves but also read the documentation for each function and use the included vignettes as examples. This eases the burden of analyzing in-house code and avoids the cost of proprietary coding software. Furthermore, the implementation of this project in R not only ensures its accessibility but also allows for integration with RAVE, another useful tool supporting open-science development of iEEG analysis.

TableContainer^2^ and Epoch^3^ also create a standardized structure for iEEG data analysis (see the following github repository for an example of how to structure data after preprocessing with RAVE^21^

https://github.com/KarasLabRAVE/RAVE_EPochDownloader). The iEEG-BIDS^18^ format exists for sharing of raw iEEG data, but these R packages allow for greater ease of analysis by addressing key barriers. They allow for storage of important metadata (e.g. patient demographics, surgical success/failure, channel information) within one single object, along with the iEEG data itself. Having this complete representation of the patient’s iEEG data makes future analysis much more streamlined, forgoing the hassle of navigating through multiple folders and files to locate relevant information. Additionally, the user can crop and transform the iEEG data while maintaining the integrity of relevant metadata, whether on a per-patient, per-electrode, or per-timepoint basis. This flexibility is key for validation of computational methods across different combinations of patients, electrodes, or time windows.

The publicly available Epoch objects on OSF^22^ also address the problem of data availability. A significant roadblock in CEZIA validation is procurement of enough properly cleaned, high-quality iEEG data to test on. The iEEG data within the OSF repository has already been reviewed, cleaned, and preprocessed, and it is ready for download and testing on EZFragility as well as other future CEZIAs.

The EZFragility package serves as the first open-science implementation of neural fragility. Despite the promising findings described in Li et al^17^, the utility of neural fragility in clinical and research settings is still limited by the complexity and inaccessibility of the algorithm. Usable source code is not provided, and there are ambiguities in the descriptions of how to calculate fragility. EZFragility^1^ addresses these problems by being publicly available on CRAN and GitHub. Its formulation as a fully documented R package allows its functions to be inspected and integrated into larger iEEG analysis pipelines. Most importantly, it is able to faithfully reproduce key figures from the original paper using the same raw data and recreates the algorithm as accurately as possible.

The EZFragility package is not an exact copy of the original neural fragility algorithm; there are important limitations on the reproducibility of the original neural fragility paper. As described previously, our fragility method differs from the original paper in the calculation of ω and automatic lambda optimization using a binary search algorithm. These were devised as solutions to computational problems we encountered while developing EZFragility: the ω was not clearly specified in prior fragility publications, and the lambda of 1e-4 as described in the original paper can result in an unstable system for some patients. Our results may also differ from the original neural fragility publication based on our independent review of channel quality, artifact identification, and seizure onset with a senior clinical epileptologist, resulting in slightly differing electrodes that were kept for analysis.

## Future work

We are continually adding more data to the OSF^22^ repository, and we are currently curating iEEG data from additional patients undergoing phase II epilepsy surgery evaluation at the University of Texas Medical Branch. We will publish them in OpenNeuro^25^ and as a data package containing Epoch objects like the one developed for this EZFragility software package demo. Current work also includes analysis on a multi-patient study of the EZFragility^17^ package results using publicly available and new data. This multi-patient data analysis will include training machine learning algorithms that allow predictions of seizure freedom after surgery based on the neural fragility computations. Finally, this ecosystem of R packages can be expanded to implement additional computational algorithms for seizure onset zone localization.

## ## acknowledgement

This work was supported by the American Society for Stereotactic and Functional Neurosurgery Research Fellowship, and Grant #BHI2024 - 16 from the Moody Brain Health Institute at UTMB.

## Notes

https://github.com/Jiefei-Wang/EZFragility

https://github.com/Jiefei-Wang/Epoch

https://github.com/Jiefei-Wang/TableContainer

### Competing Interest Statement

The authors have declared no competing interest.

